# Eyeblink conditioning performance and brain-wide *C-fos* expression in male and female mice

**DOI:** 10.1101/2021.10.15.464518

**Authors:** Maria Roa Oyaga, Ines Serra, Devika Kurup, Sebastiaan K.E. Koekkoek, Aleksandra Badura

## Abstract

The functional and molecular sources of behavioral variability in mice are not fully understood. As a consequence, the predominant use of male mice has become a standard in animal research, under the assumption that males are less variable than females. Similarly, to homogenize genetic background, neuroscience studies have almost exclusively used the C57BL/6 (B6) strain. Here, we examined individual differences in performance in the context of associative learning. We performed delayed eyeblink conditioning while recording locomotor activity in mice from both sexes in two strains (B6 and B6CBAF1). Further, we used a C-FOS immunostaining approach to explore brain areas involved in eyeblink conditioning across subjects and correlate them with behavioral performance. We found that B6 male and female mice show comparable variability in this task and that females reach higher learning scores. We found a strong positive correlation across sexes between learning scores and voluntary locomotion. C-FOS immunostainings revealed positive correlations between C-FOS positive cell density and learning in the cerebellar cortex, as well as multiple previously unreported extra-cerebellar areas. We found consistent and comparable correlations in eyeblink performance and *C-fos* expression in B6 and B6CBAF1 females and males. Taken together, we show that differences in motor behavior and activity across brain areas correlate with learning scores during eyeblink conditioning across strains and sexes.

## Introduction

For several decades, female mice have been considerably under-investigated in neuroscience due to the presumption that hormonal fluctuations caused by the estrous cycle might introduce non-comparable variability across sexes (Meziane et al., 2007). However, meta-analyses of rat and mice studies show that females and males exhibit comparable variability across behavioral, morphological and physiological traits, and that for most traits, female estrous cycle does not need to be considered (Simpson and Kelly, 2012; Becker et al., 2016). Notwithstanding, female and male mice show sex-specific strategies in locomotion adaptation, reward learning, and spatial orientation and learning (Konhilas et al., 2004; Bettis and Jacobs, 2009; Hendershott et al., 2016; Grissom et al., 2018; Prawira, 2019).

The development of inbred mouse strains was also intended to decrease the variability between animals and increase the power of studies (Festing, 1999). However, the current golden standard to keep mice exclusively on the C57BL/6 (B6) strain, limits the generalization of findings (Rivera and Tessarollo, 2008; Sittig et al., 2016). When it comes to inter-strain variability, behavioral differences have been reported, yet, it has not been systematically investigated across commonly used paradigms (Faure et al., 2017; Arnold and Newland, 2018).

Although some components of murine behavioral variability have been extensively studied, a number of variables that could alter learning are largely unexplored (Pfaff, 2001; Bucán and Abel, 2002; Tye et al., 2011; Leung and Jia, 2016). Hence, gaining a deeper understanding of the sources of behavioral variability could give us indications on how to interpret data and ensure better reproducibility across laboratories.

In order to study how sex and strain influence mouse behavior and brain activity, a reliable and controlled paradigm is needed. We investigated this in the context of delay eyeblink conditioning. Eyeblink conditioning is a cerebellar-dependent associative learning paradigm, in which an initially neutral, conditioned stimulus (CS, a flashing light), becomes predictive of an unconditioned stimulus (US, an air-puff to the cornea), which elicits a blink. The paradigm consists of pairing the CS with the US and, over time, an association is formed where blinking is triggered by the CS alone. The newly learned association is called conditioned response (CR) (Gormezano et al., 1962).

There is limited knowledge on sex-related performance differences in eyeblink conditioning. In human delay eyeblink studies, girls show more CRs in the first five days of learning compared to boys, and women show a continuous increase in CRs compared to men (Löwgren et al., 2017). In rats, stress seems to enhance delay eyeblink conditioning in males but hinders learning in females (Wood and Shors, 1998). However, in rabbits, males and females show similar conditioning profiles but females seem to adapt faster to stress (Schreurs et al., 2018). Finally, in mice, females show increased CRs compared to males in the first five days of learning trace conditioning (Rapp et al., 2021), a different form of eyeblink conditioning where a CS and US are separated in time. Comparison of performance differences in eyeblink conditioning in different strains is currently lacking.

At the circuit level, studies have shown that the association between the stimuli during delay eyeblink conditioning most likely relies on the cerebellum. Here, the CS signals coming from the pons and the CS information from the inferior olive via climbing fibers are precisely timed and processed (Heiney et al., 2014a; ten Brinke et al., 2015). Several cerebellar areas modulating eyeblink conditioning have been identified in mice; lobule VI in the vermal region and crus 1 and simplex in the hemispheric region (Heiney, Kim, et al., 2014; Gao et al., 2016). Inactivation of lobule VI and crus 1 during development causes deficits in learning, indicating a crucial role in eyeblink conditioning (Badura et al., 2018). The CR signal leaves the cerebellum via the interposed nucleus, which ultimately connects to the muscles controlling the eyeblink reflex (Gao et al., 2016; ten Brinke et al., 2017). Beyond the pontocerebellar and olivocerebellar systems, little is known about the potential involvement of other brain areas in eyeblink conditioning (Boele et al., 2010; Ruigrok, 2011; D’Angelo et al., 2016; Kratochwil et al., 2017). The amygdala has been proposed to have a role in associative learning, given its implication in fear conditioning and arousal (Lee and Kim, 2004). Specifically, lesions in the amygdala during the first days of training highly impair learning, while lesions in later stages do not appear to affect learning (Lee and Kim, 2004).

Our understanding of behavioral variability and engagement of different brain regions is finally limited by the practise of outliers removal - animals that deviate from the group mean and do not reach proficient learning scores are commonly dropped from further analysis (Osborne and Overbay, 2004; Rousselet and Pernet, 2012; Fonnesu and Kuczewski, 2019). Possible mechanisms underlying performance differences could be arousal levels and locomotor activity, which both influence cortical function (McGinley et al., 2015; Vinck et al., 2015; Williamson et al., 2015), or stress levels which can affect neuronal firing in the deep cerebellar nuclei (DCN) and hippocampus (Joëls, 2009; Schneider et al., 2013). In the cerebellum, although locomotion modulates activity in the cortex, the relevance of this modulation is still not fully understood (Ozden et al., 2012; Hoogland et al., 2015; Powell et al., 2015). During eyeblink conditioning, imposed locomotor activity enhances learning by increased activation of the mossy fiber pathway to the cerebellar cortex (Albergaria et al., 2018).

Here, we investigated the effect of sex in behavioral variability by employing eyeblink conditioning to quantify performance differences in B6 mice and B6CBAF1 mice. Furthermore, we explored the engagement of brain regions that may have a modulatory role in eyeblink conditioning by utilizing *C-fos* expression as a proxy for neural activity during learning in both of those strains. We found that male and female mice of both B6 and B6CBAF1 strains showed comparable variability in the delay eyeblink conditioning. However, females reached higher learning scores. Further, we found a strong positive correlation across sexes between learning scores and voluntary locomotion in the B6 mice. C-FOS immunostaining revealed positive correlations between C-FOS positive cell density and learning in the cerebellar cortex, as well as multiple previously unreported extra-cerebellar areas.

## Results

### B6 female and male mice show comparable variability in eyeblink conditioning and females reach higher learning scores

To study differences in learning profiles between sexes, we performed delayed eyeblink conditioning experiments with B6 females (n = 14) and males (n = 14). First, we habituated the animals to the set-up for increasing periods of time over five days to decrease anxiety levels and optimize training. Next, we subjected mice to a 5-day training paradigm in order to capture behavioral variability in the first stages of learning. This training length was selected considering that animals show the most variability within the first days of acquisition, start showing reliable CRs in day four/five and eventually plateau during the last 5 days of training (Heiney et al., 2014b; Giovannucci et al., 2017).

The blue LED light (CS) was triggered 250 ms prior to the puff to the cornea (US) in paired trials and the two stimuli co-terminated (**Fig. 1A**). Sessions consisted of 20 blocks of 12 trails each (1 US only, 11 paired and 1 CS only). Mice learned the association between the stimuli progressively and developed a gradually increasing conditioned response (CR) (**Fig 1A, 1B**). Males and females had comparable learning profiles, and the variances during training sessions were not significantly different between sexes (F-test for two sample variances in CR amplitude of paired trials, F = 3.66, *p* = 0.11). In CS only trials, females showed a slight increase in CR percentage on session four, that culminated with a 60% CR responses in session five opposed to 40% in males (two-way ANOVA repeated measures for sex and sessions: sex effect: F(1,26) = 1.461, *p* = 0.24, interaction sex and session: F(4,104) = 4.01, *p* = 0.02, Cohen’s d session five: 0.88) (**Fig. 1C**). The CR amplitude (measured as the response normalized to UR max amplitude = 1) during CS trials was significantly higher in females compared to males (two-way ANOVA repeated measures for sex and sessions: sex effect: F(1,26) = 6.109, *p* = 0.02, interaction sex and session: F(4,104) = 5.12, *p* = 0.0008, Cohen’s d session five: 0.93) (**Fig. 1D**). On the last session of training, females reached an average amplitude of 0.55 while males reached an average of 0.33 (**Fig. 1E**). Overall, these results show that male and female mice show comparable variance in eyeblink conditioning, but females reach higher learning scores in a 5-day training paradigm.

**Figure 1:**
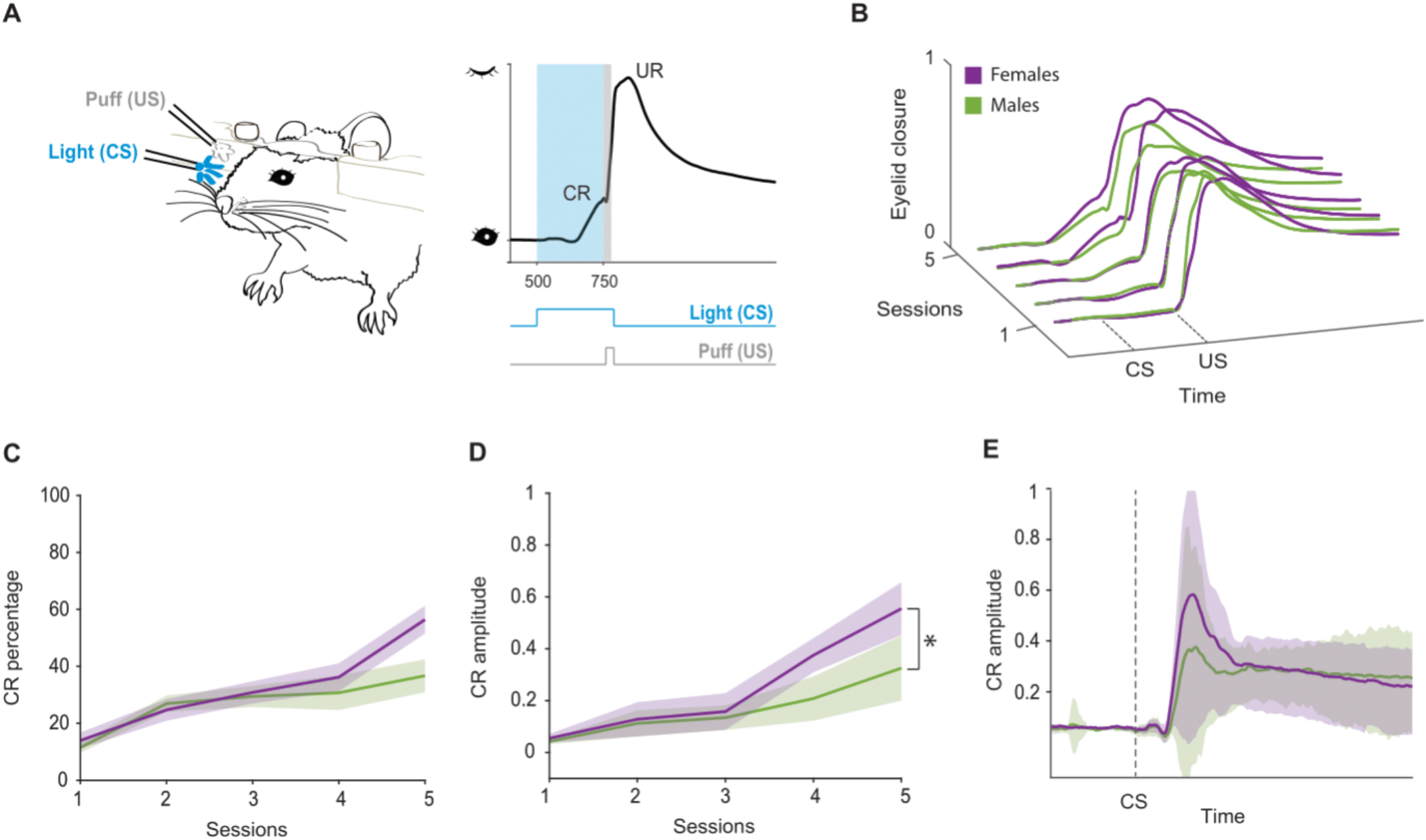
B6 female and male mice show comparable variability in eyeblink conditioning and females reach higher learning scores. A) Experimental setup. Mouse with implanted headplate is head-fixed on top of a freely rotating wheel. A blue light (conditioned stimulus, CS) is presented 250 ms before a puff (unconditioned stimulus, US) to the same eye. In a trained mouse, the CS produces an anticipatory eyelid closure (conditioned response, CR) followed by a blink reflex triggered by the US (unconditioned response, UR). B) Paired trails average traces in females and males over training sessions. The CR progressively develops due to the CS-US paring. C) CR percentage in CS only trials over training sessions. Purple: females, green: males. Shaded area: sem. D) CR amplitude in CS only trials over training sessions (two-way ANOVA for sex and sessions: sex effect: F(1,26) = 6.109, p = 0.02) Shaded area: sem. E) Average response in CS only trials in the last session of training. Purple: females (n=14), green: males (n=14). Shaded area: std.

### Learning scores correlate with spontaneous locomotor activity

We next asked whether mice show behavioral differences in voluntary locomotor activity during training, as it has been shown that imposed locomotion affects eyeblink performance (Albergaria et al., 2018). We investigated whether higher learning scores would correlate with higher spontaneous locomotor activity. For this purpose, we added infrared cameras to the eyeblink setups to record body movements during eyeblink sessions. The cameras were placed at the right back corner of the box, allowing a wide recording angle to capture whole body movements (**Fig. 2A**). We recorded videos of full training sessions for each mouse, which were later analyzed offline. The head bar height (**Fig. 1A**) was adjusted accordingly to ensure that every mouse could move comfortably on the wheel.

**Figure 2:**
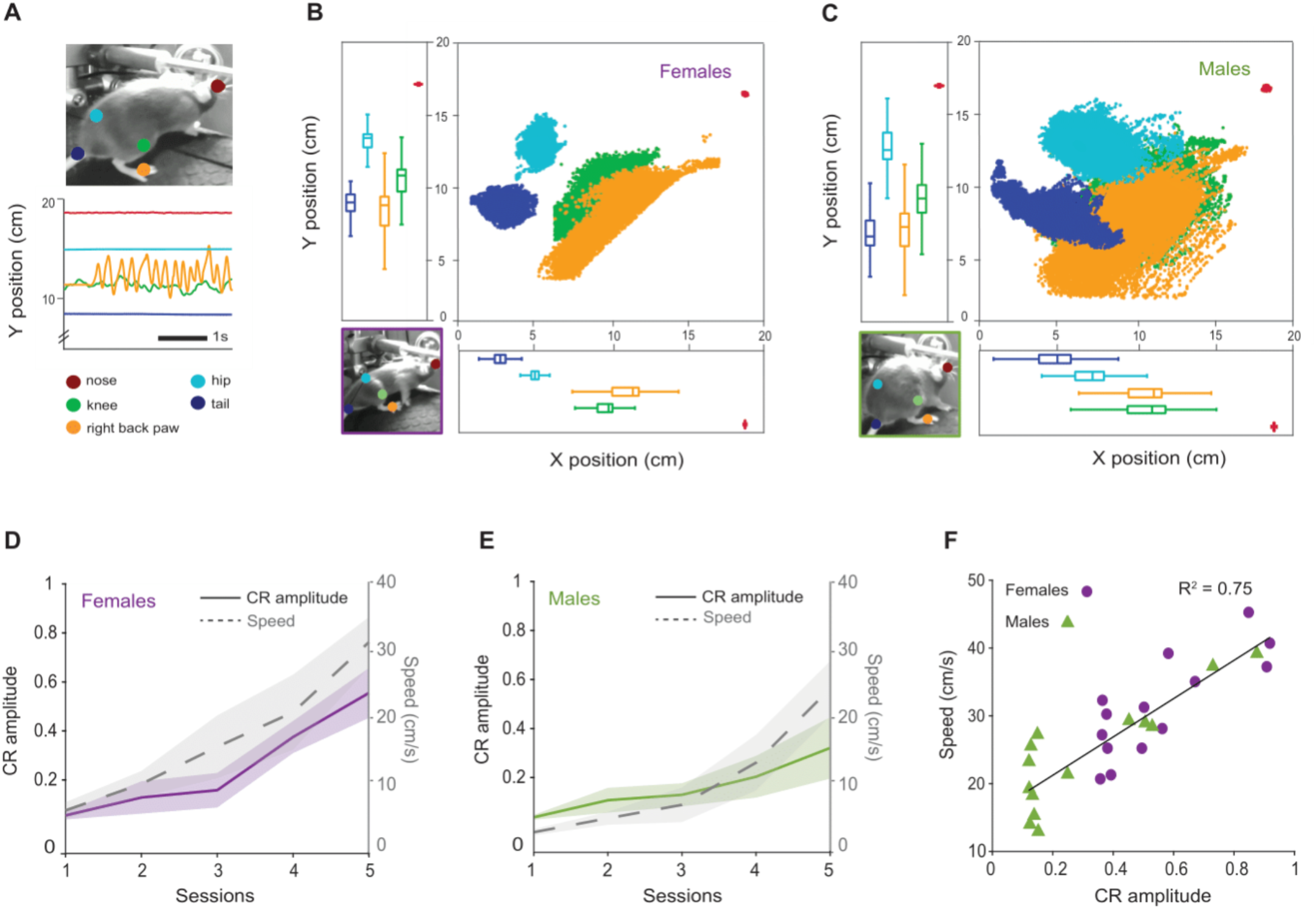
Learning scores correlate with spontaneous locomotor activity. A) Top: Example view of tracking with DeepLabCut. Bottom: Example tracking traces (Y position change). B) Scatter plot and boxplots of each body part (females, n = 14). C) Scatter plot and boxplots of each body part (males, n = 14). D and E) CR amplitude and speed of the right back paw over training sessions. Purple: females (n = 14), green: males (n = 14). Shaded area: sem. Speed: two-way ANOVA for sex and sessions: sex effect: F(1,26) = 12.17, p = 0.0017. F) Positive correlation between CR amplitude and speed of the right back paw on the last session of training (linear regression: R^2^ = 0.75, p = 0.002).

To track different body parts and get a meaningful movement output, we used DeepLabCut (DLC), a software for automated animal pose tracking (Mathis et al., 2018) (Materials and Methods; *Movement analysis*). This approach allows movement tracking without utilizing physical markers on the body that can hinder natural movement. We tracked 5 body parts: tail base, hip, knee, right back paw and nose (**Fig. 2A**). Animals were head-fixed on top of the wheel, hence, Y position was similar between both sexes (**Fig 2. B, C**). We observed more fluctuation in the hip and tail along the X axis in males, which might indicate a tendency to rotate their body axis from side to side (**Fig 2. B,C**). We selected speed of the right back paw as a proxy for general locomotion behavior on further analysis.

Animals increased their speed on the wheel during training, and both females and males had comparable variances (F-test for two sample variances in speed: F = 0.58, *p* = 0.305). We found that females moved significantly faster than males during learning, reaching an average speed of 31 cm/s compared to males that reached 24 cm/s (two-way ANOVA repeated measures for sex and sessions: sex effect: F(1,26) = 12.17, *p* = 0.0017, Cohen’s d session five: 1.07) (**Fig. 2D, E**). Because we observed a similar trend between CR amplitude and running speed across sexes, we performed a linear regression between speed of the right back paw and CR amplitude in the last session. This showed a clear correlation between the variables (R^2^ = 0.75, *p* = 0.002) (**Fig.2 F**). These results reveal that mice that spontaneously move faster on the wheel, reach higher learning scores in eyeblink conditioning.

### Learning scores correlate with C-fos expression

To explore if differences in eyeblink performance correlate with differences in brain activity, we performed C-FOS immunostainings following the last training session. *C-fos* is an immediate early expressed gene, a family of transcription factors that is expressed shortly after a neuron has depolarized. Because of its precise time window of expression, it is widely used as an activity marker (Chung, 2015). Evidence suggests that C-FOS protein greatly increases after exposure to novel objects, surroundings or stimuli, while continuous, long-term exposure to persistent stimuli returns *C-fos* expression to basal levels (Joo et al., 2015; Gallo et al., 2018; Bernstein et al., 2019). Besides our aim to capture the variability in the first stages of learning, the *C-fos* expression due to novelty was another reason to choose a 5-day training paradigm instead of the standard 10-day acquisition training.

We imaged sections of whole brains with a fluorescent microscope and developed an image analysis workflow to quantify C-FOS positive neurons and identify their location within the hierarchical structure of the Allen Brian Atlas (cerebrum, brainstem and cerebellum). (**Supp. Fig. 1**).

To identify regions potentially involved in associative learning, we selected mice that showed a CR amplitude of 0.4 or higher in CS only trials (n = 14; 5 B6 males, 9 B6 females). These mice are further referred as “learners”. We performed Kendall’s correlation (non-parametric rank order regression) between density of C-FOS positive cells and CR amplitude on the last session. In the cerebellum, C-FOS labeling was localized in the granule cell layer, which we confirmed with colocalization with GABAα6, a granule cell specific marker (**Fig 3A, Supp Fig. 2, Supp Fig. 3 A, B**). In the cerebellar hemispheric regions, crus 1 and simplex had a significant correlation between C-FOS cell density and CR amplitude (crus 1: tau = 0.42, *p* = 0.042, simplex: tau = 0.52, *p* = 0.009). In the cerebellar vermis, lobule VI also had a significant correlation and the highest Tau (lobule VI: tau = 0.8, *p* = 0.009) (**Fig. 3B**). In the brain stem, we found a significant correlation in the red nucleus, the facial nucleus, the inferior olive and the pontine nuclei (Red nucleus: tau = 0.43, *p* = 0.041, Facial nucleus: tau = 0.57, *p* = 0.006, inferior olive: tau = 0.76, *p* = 0.0008, pontine nuclei: tau = 0.48, *p* = 0.021) (**Fig. 3E, Supp Fig. 2**). We found a positive correlation in the visual, motor and somatosensory cortices, and the amygdala (visual cortex: tau = 0.51, *p* = 0.013, motor cortex: tau = 0.69, *p* = 0.0003, somatosensory cortex: 0.54, *p* = 0.007, amygdala: tau = 0.63, *p =* 0.001) (**Fig. 3H, Supp Fig. 2**). Finally, to understand whether there was a certain layer specificity in these cortices, we performed two double immunostainings with C-FOS; CUX1, a marker for upper cortical layers (II-IV) and CTIP2, for lower cortical layers (V-VI). Although we detected C-FOS positive cells in all layers of the cortex, we observed a higher colocalization of CUX1 and C-FOS compared to CTIP2 and C-FOS (**Supp Fig. 3 C-F**).

**Figure 3:**
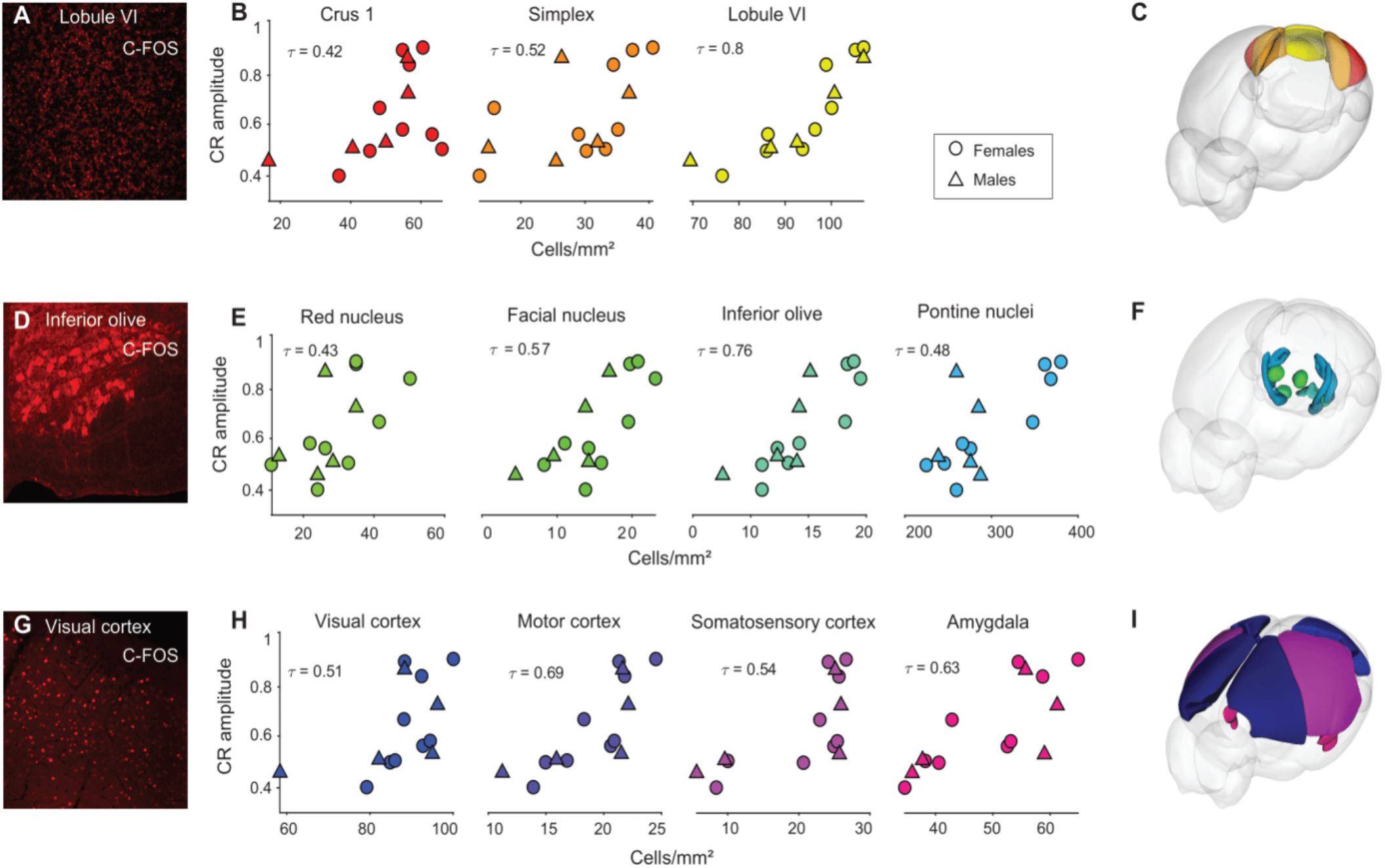
Learning scores correlate with *C-fos* expression. **A**) C-FOS positive granule cells in lobule VI in the cerebellum. **B**) Cerebellar areas with significant positive correlation between C-FOS positive cell density and CR amplitude (crus 1: tau = 0.42, *p* = 0.042, simplex: tau = 0.52, *p* = 0.009, lobule VI: tau = 0.8, *p* = 0.009). **C**) 3D model with significant areas highlighted. **D**) C-FOS positive cells in the inferior olive. **E**) Brainstem areas with significant positive correlation between C-FOS positive cell density and CR amplitude (red nucleus: tau = 0.43, *p* = 0.041, facial nucleus: tau =0 .57, *p* = 0.006, inferior olive: tau = 0.76, *p* = 0.0008, pontine nuclei: tau = 0.48, *p* = 0.021). **F**) 3D model with significant areas highlighted. **G**) C-FOS positive cells in the visual cortex. **H**) Cortical areas with significant positive correlation between C-FOS positive cell density and CR amplitude (visual cortex: tau = 0.51, *p* = 0.013, motor cortex: tau = 0.69, *p* = 0.0003, somatosensory cortex: 0.54, *p* = 0.007, amygdala: tau = 0.63, *p*: 0.001). **I**) 3D model with significant areas highlighted.

Together, these results confirm the previously reported areas associated with eyeblink conditioning within the olivo-cerebellar and ponto-cerebellar systems (Ruigrok, 2011; D’Angelo et al., 2016) and suggest that other areas might be involved in this learning task.

### Pseudoconditioned mice show lower C-fos expression

To ensure an appropriate control for the quantification of C-FOS positive cells, we included pseudoconditioned mice (n = 2 B6 males, 2 B6 females) in our experimental design. These mice went through the same experimental steps as the conditioned mice, with the only exception that they were not trained with paired CS-US trials. Instead, we exposed them to a protocol with CS and US only trials, keeping the same structure and duration as the conditioned protocol. Pseudoconditioned mice did not acquire an association given that there was no substrate for learning (absence of CS-US pairing) and exhibited a slight increase in locomotion speed over training sessions (**Fig. 4A**). We used the same analysis method to quantify the density of C-FOS positive cells in pseudoconditioned mice. Overall, we observed lower *C-fos* expression in pseudoconditioned mice compared to learners (**Fig. 4B**). We compared the C-FOS density between pseudoconditioned mice and learner mice in the areas where we had found a positive significant correlation between CR amplitude and C-FOS density (**Fig. 3**). Learners had a higher C-FOS density median compared to pseudoconditioned mice (**Fig. 4C**). We found a significant difference between the groups in each one of these areas (Two-tailed Mann-Whitney U test between pseudoconditioned and learner mice; crus 1: *p* = 0.00008, simplex: *p* = 0.00008, lobule VI: *p* = 0.00008, red nucleus: *p* = 0.00008, facial nucleus *p* = 0.01, inferior olive: *p* = 0.00008, pontine nucleus: *p* = 0.00008, visual cortex: *p* = 0.00008, motor cortex: *p* = 0.0002, somatosensory cortex: *p* = 0.03, amygdala: *p* = 0.001) (**Fig. 4D**). This confirms that the observed increased *C-fos* expression in learner mice is due to the CS-US pairing.

**Figure 4:**
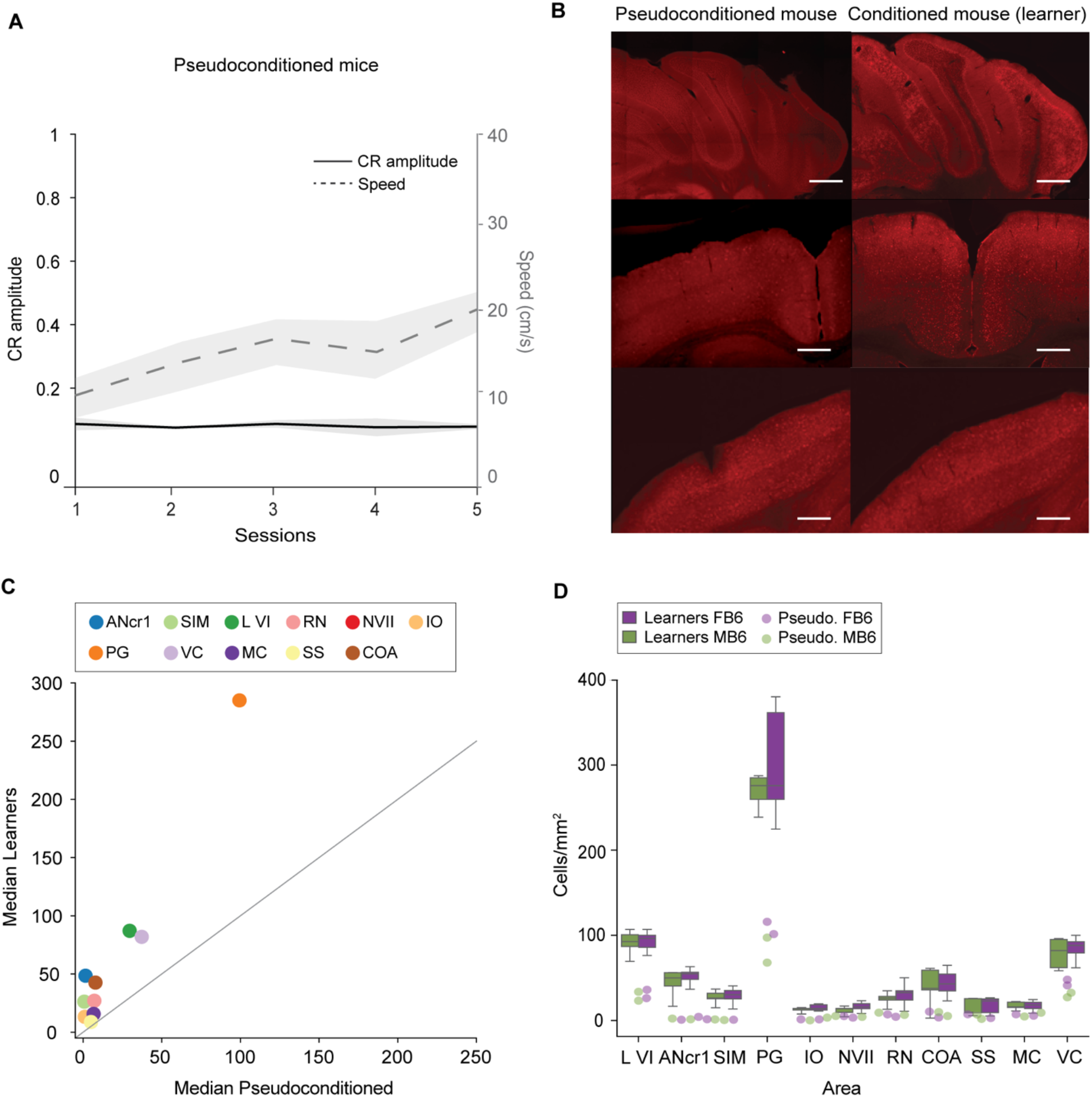
Pseudoconditioned mice show lower *C-fos* expression compared to learners. **A)** Change in CR amplitude and Speed of the right back paw over training sessions. CR amplitude left Y axis in black, Speed, in the right Y axis in grey. **B**) Immunofluorescent images, C-FOS positive cells. Top to bottom: cerebellar cortex, motor cortex, somatosensory cortex. Scale bar: 100 μm. **C)** Median pseudoconditioned mice against median learners, unity line in grey. **D**) Boxplots depict C-FOS density in learners (n = 9 females, 5 males), dots are pseudoconditioned mice (n = 2 females, 2 males). Green males, purple females. Brain area acronyms from the Allen Brain Atlas: SIM: simplex, ANcr1: crus 1, L VI: lobule VI, RN: red nucleus, NVII: facial nucleus, IO: inferior olive, PG: pontine nucleus, VC: visual cortex, MC: motor cortex, SS: somatosensory cortex, COA: amygdala.

### Correlation between learning scores and C-fos expression is consistent in B6CBAF1 strain

Given the evidence of the possible unwanted effects of highly inbred mouse strains like B6 in replicability and reproducibility (Åhlgren and Voikar, 2019), we wanted to investigate inter-strain variability in associative learning. We asked whether the results obtained in B6 mice would be consistent in a different mouse strain. For this purpose, we performed eyeblink conditioning together with C-FOS immunostainings in B6CBAF1 mice, which are the F1 hybrids of B6 and CBA strains. Hybrid mice are used due to their hybrid vigor, the robustness and health gained from a high degree of heterozygosity (Wolfer et al., 2002). B6CBAF1 mice have significantly less retinal degeneration and hearing loss compared to B6 mice, which makes them an appropriate candidate for visual and auditory experiments (Erway et al., 1996; Milon et al., 2018; Ohlemiller, 2019).

B6CBAF1 mice learned the association between the stimuli and gradually formed CRs. We observed a trend indicating similar sex differences between B6CBAF1 mice and B6. Females reached 53% CR percentage compared to 35% in males (two-way ANOVA repeated measures for sex and sessions: sex effect: F(1,14) = 2.55, *p* = 0.237, interaction sex and session: F(4,104) = 3.01, *p* = 0.021, Cohen’s d session five: 0.86) (**Fig 5. A**). When it comes to the amplitude of these responses, BFCBAF1 females showed a trend towards slightly higher CR amplitude over training sessions compared to males (two-way ANOVA repeated measures for sex and sessions: sex effect: F(1,14) = 2.55, *p* = 0.132, Cohen’s d session five: 0.84) (**Fig 5. B**).

**Figure 5:**
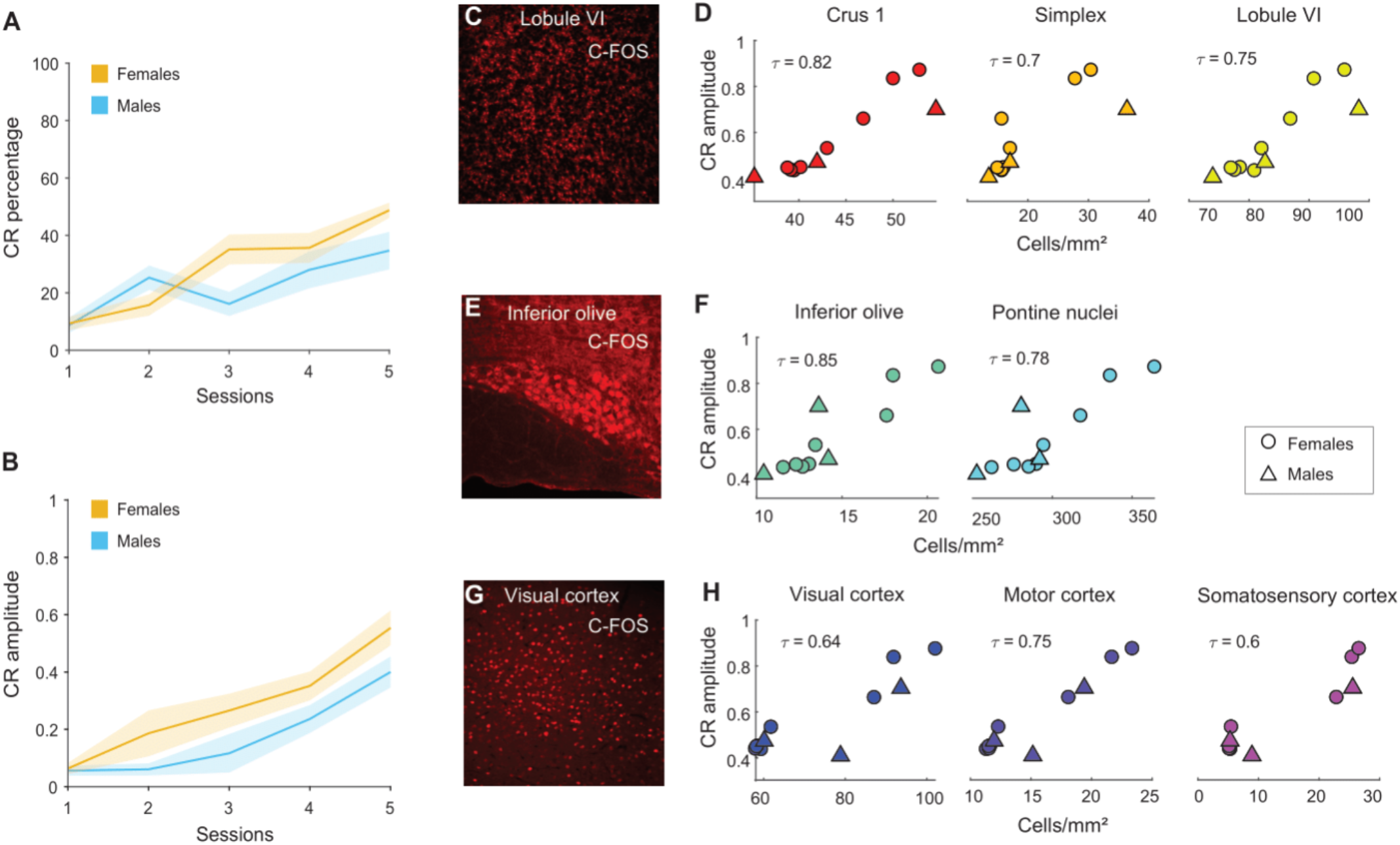
Correlation between learning scores and *C-fos* expression is consistent in B6CBAF1 mice. **A**) CR percentage in CS only trials over training sessions. Yellow: females (n = 9), cyan: males (n = 7). Shaded area: sem. **B**) CR amplitude in CS only trials over training sessions. Shaded area: sem **C**) C-FOS positive cells in the lobule VI. **D**) Cerebellar areas with significant positive correlation between C-FOS positive cell density and CR amplitude (crus 1: tau = 0.82, *p* = 0.0001, simplex: tau = 0.7, *p* = 0.005, lobule VI: tau = 0.75, *p =* 0.0007). **E**) C-FOS positive cells in the inferior olive. **F**) Brainstem areas with significant positive correlation between C-FOS positive cell density and CR amplitude (inferior olive: tau = 0.85, *p =* 0.0004, pontine nuclei: tau = 0.78, *p =* 0.0003). **G**) C-FOS positive cells in the visual cortex. **H**) Cortical areas with significant positive correlation between C-FOS positive cell density and CR amplitude (visual cortex: tau = 0.64, *p* = 0.0057, motor cortex: tau = 0.75, *p* = 0.0008, somatosensory cortex: tau = 0.6, *p =* 0.009).

We followed the same analysis pipeline to quantify *C-fos* expression in brain slices of B6CBAF1 mice after eyeblink conditioning. Next, we selected mice that showed a CR amplitude of 0.4 or higher in CS only trials (n =11; 3 males, 8 females) and performed Kendall’s correlation between density of C-FOS positive cells and CR amplitude on the last training session. The granule cell layer also contained C-FOS labelling, and crus 1, the simplex and lobule VI were found to have a significant positive correlation (crus 1: tau = 0.82, *p* = 0.0001, simplex: tau = 0.7, *p* = 0.005, lobule VI: tau = 0.75, *p* = 0.0007) (**Fig 5. C, D, Supp Fig. 5**). In the hindbrain, the correlation between C-FOS cells and learning was also significant in the inferior olive and the pontine nuclei (inferior olive: tau = 0.85, *p* = 0.0004, pontine nuclei: tau = 0.78, *p* = 0.0003) (**Fig 5. E, F, Supp Fig. 5**). Additionally, the visual, motor and somatosensory cortices showed significant positive correlations (visual cortex: tau = 0.64, *p* = 0.0057, motor cortex: tau = 0.75 *p* = 0.0008, somatosensory cortex: tau = 0.6, *p* = 0.009) (**Fig 5. G, H, Supp Fig. 5**). These results show that most of the areas where we saw a correlation between learning and C-FOS density in B6 mice are also found in B6CBAF1 mice, which strengthens the idea that these areas are active during eyeblink conditioning.

### Variability between sexes and strains

To further understand inter-strain and inter-sex variability in our dataset, we calculated the coefficient of variance (CV, standard deviation/mean) for each of the variables that we quantify in both B6 and B6CBAF1 mice. We selected the 14 learners (CR amplitude on session 5 > 0.4) B6 mice (n = 5 males, 9 females) and the 11 learners B6CBAF1 mice (n = 3 males, 8 females) and grouped them by strain and sex (**Fig. 6**). For each group, we calculated the CV for the common variables acquired and previously reported, which can be grouped in two main categories: eyeblink performance and *C-fos* expression. Eyeblink performance includes CR amplitude and percentage. *C-fos* expression includes the density of C-FOS positive cells in the brain areas where we have found a positive significant correlation across both strains: crus 1, simplex, lobule VI, inferior olive, pontine nuclei, and the visual, motor and somatosensory cortices. When comparing strains, we observed that the variances for each variable were similar, with the exception of the somatosensory cortex, where B6CBAF1 mice seem to be more variable compared to B6 (**Fig. 6A**). B6 female and male mice had similar CVs for most of the variables, although in crus 1 and the somatosensory cortex males seem to have a slightly higher CV (**Fig. 6B**). However, this could be due to the difference in sample size. We observed something similar between B6CBAF1 female and male mice; males showed slightly higher CV in C-FOS density in the simplex (**Fig. 6C**). Additionally, B6CBAF1 mice had the highest CV in the somatosensory cortex.

**Figure 6:**
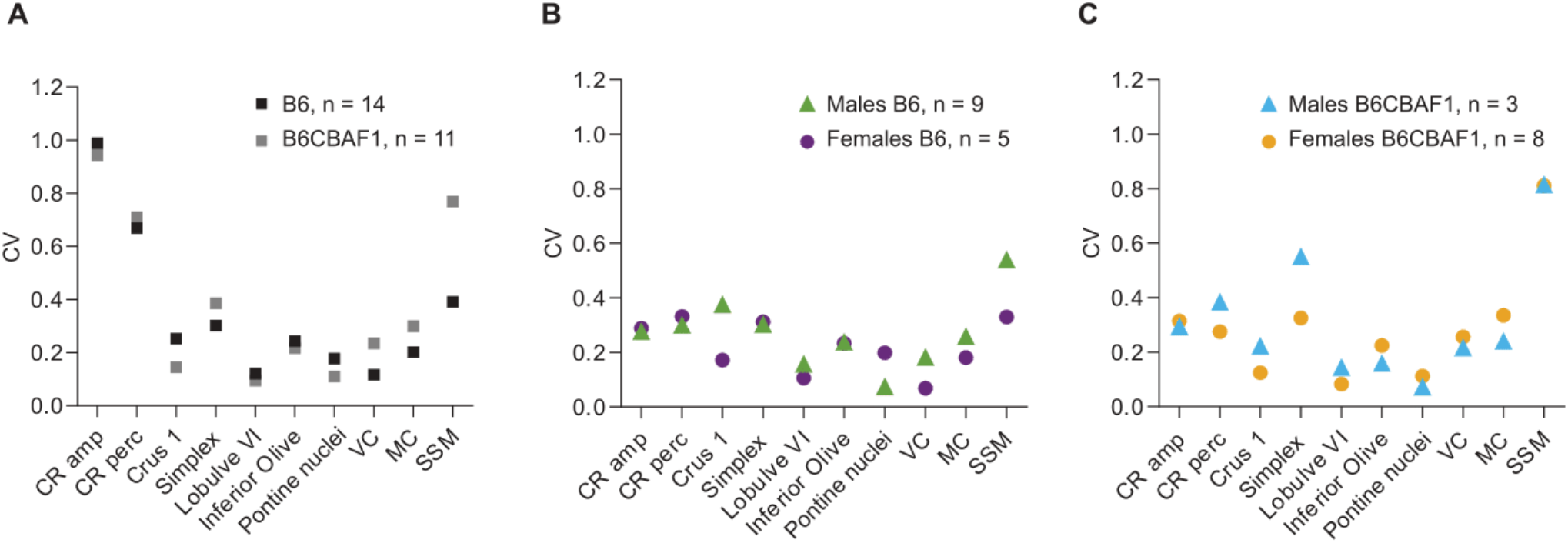
Strain and sex variability. Learners were selected if CR amplitude on session 5 > 0.4. CV = STD/mean. **A**) For strains: B6, n = 14, B6CBAF1, n = 11. **B**) For B6 mice: males B6, n = 9, females B6, n = 5. **C**) For B6CBAF1 mice: males, n = 3, females, n = 8. VC: visual cortex; MC: motor cortex; SSM: somatosensory cortex; CV: Coefficients of variation.

## Discussion

Understanding behavioral variability in the context of neuroscience research is a challenge. We are still far from fully comperhending how factors like sex and strain give rise to differences in behavior.

Here we addressed this question by making use of a well-known learning paradigm to study behavioral variability. We found that B6 female and male mice showed comparable variance in delay eyeblink conditioning and locomotion while being head-fixed on a rotating wheel. The variance within these behaviors was not different between sexes, yet females reached higher learning scores and running speeds within five days of training. Importantly, we found a robust correlation between learning scores and running speed which is consistent across sexes. In a similar way, we found that enriched *C-fos* expression across several brain areas positively correlates with learning, which suggests the involvement of these regions in eyeblink conditioning. Finally, we observed similar results in a hybrid mouse strain (B6CBAF1).

### Sex, strain and behavior: comparable variability but difference in performance

Opposed to what is sometimes assumed in behavioral science, we observed that sexes show comparable variability. However, we found differences in performance during eyeblink conditioning and locomotion. Common behaviors like duration of running vary between female and male mice in the wild (Lightfoot et al., 2004; Goh & Ladiges, 2015). Behaviors widely assessed in research such as fear conditioning and navigation on the Morris water maze also show differences depending on sex (Roof & Stein, 1999; Keeley et al., 2013; Yang et al., 2013; Gruene, Flick, et al., 2015). In the context of cerebellar-dependent learning, evidence shows that estradiol increases the density of parallel fiber to Purkinje cell synapse and induces long-term potentiation, which improves memory formation (Andreescu et al., 2007). In trace eyeblink conditioning, both sexes reach similar learning scores but females show significantly higher CR percentage compared to males in the first five days of learning, which is in line with our findings (Rapp et al., 2021). Considering that adapting motor reflexes is a highly conserved behavior, it is logical that both sexes reach similar asymptotic learning scores in longer paradigms. However, these findings together with our results suggest that females exhibit faster learning rates during the first stages of learning.

Therefore, we postulate that using females could reduce the training time to achieve desired scores, which could be advantageous for certain experiments, particularly time-sensitive ones, such as calcium imaging or electrophysiological measurements. In addition, our results show that male and female mice have similar variability, which indicates that females can be included in studies without taking into account the estrous cycle phase. In general, utilizing both sexes would reduce the overall number of animals used in research, and increase the relevance and generalization of scientific findings.

A common experimental setting in neuroscience involves head-fixing awake mice and placing them on a freely moving wheel. A recent study investigated sex differences in head-fixed running behavior and found that female mice ran forward naturally within the first two days, while males took seven days to progressively learn to only run forward (Prawira, 2019). In our experiments, the differences in learning scores between sexes were strongly correlated to the changes in locomotor activity on the wheel. Our results show that the previously reported correlation between imposed locomotor activity and learning scores (Albergaria et al., 2018) persists when mice can initiate locomotion voluntarily. This suggests that spontaneous locomotion might facilitate associative learning and could be predictive of learning scores.

Finally, we have found that, besides moving slower, males tended to have a tilted position on the wheel compared to females. These differences in body position could be partially caused by differences in stress levels that, at the same time, could affect learning rates. It is known that stress plays an important role in modulating neural activity in the hippocampus. Corticosterone - among other stress hormones – increases CA1/CA3 firing rates shortly after a stressful period and induces molecular cascades that enhance calcium influx, which disrupts hippocampal function (Joëls, 2009). Similar mechanisms have been described in the cerebellum; calcium-based excitability in the DCN is altered in animals with higher levels of corticosterone evoked by shipping stress (Schneider et al., 2013).

### Associative learning networks

Our results show that the expression of *C-fos* in the cerebellar cortex, following delay eyeblink conditioning, is localized to the granule cell layer. This is expected, given that multiple forms of plasticity have been shown within the synapses in this layer. For example, the mossy fiber-granule cell synapse undergoes both long-term potentiation and long-term depression (Gao et al., 2012), and evidence has shown that granule cell activity adapts over time during eyeblink conditioning (Giovannucci et al., 2017) and other types of learning (Knogler et al., 2017; Wagner et al., 2017). In addition, induction of LTP by theta-burst stimulation in acute cerebellar slices activates cAMP-responsive element binding protein (CREB) cascade which, in turn, activates *C-fos* expression (Gandolfi et al., 2017). Our results are consistent with these findings and, overall, they provide evidence on how plasticity, at the input level in the cerebellar cortex, can evoke transcriptional processes that contribute to learning consolidation. The strongest correlations between *C-fos* expression and CR amplitude within the cerebellum were observed in simplex, lobule VI, and crus 1, which is consistent with the “eyeblink region”, but expands beyond the small area usually recorded using electrophysiological approaches (Heiney et al., 2014a; ten Brinke et al., 2015). Strong *C-fos* expression in crus 1 supports our previous findings showing the importance of this lobule in eyeblink conditioning (Badura et al., 2018).

Outside of the cerebellum, we identified several brain areas that could play a role in eyeblink conditioning. At the brainstem level, we found a relation between high learning scores and *C-fos* expression in the pons, inferior olive, red nucleus and facial nucleus. High activity in the red nucleus and facial nucleus is to be expected, given that these two nuclei, together with the oculomotor nucleus, execute the blink. The inferior olive and the pontine nuclei relay the US and CS information to the cerebellar cortex, respectively. During early training sessions, the US is a highly aversive stimulus, which makes it comparatively more salient than the CS signal. Hence, one would expect increased activity in the pons relative to the inferior olive. However, in later learning stages (when animals have consolidated the association), the CS is predictive of the US, which would increase the activity in the inferior olive relative to the pons. We found a correlation between learning scores and C-FOS positive cells in both the pons and the inferior olive, which could indicate an intermediate stage of learning, where animals have learned the association but the US information is still relevant.

Moreover, we found higher *C-fos* expression in the visual, motor and somatosensory cortices in mice with higher learning scores. Processing in these cortices could facilitate the CS to become more salient and ultimately predict the US. The somatosensory cortex projects to the lateral amygdala which, in turn, projects to the central amygdala, to ultimately contact the pons. The high *C-fos* expression found in the amygdala points towards a two stage conditioning model; where the amygdala would have an initial role with arousal as a salient feature, and a second phase where the cerebellum would take over to form precisely-timed CRs (Boele et al., 2010). In the motor cortex, higher C-FOS levels in high performing mice might be due to locomotor activity rather than learning itself. However, as mentioned above, this could play a role in learning either by directly affecting cerebellar input or indirectly as arousal.

Finally, we found an enrichment of *C-fos* expression in upper cortical layers (II, III and IV), especially in the visual and somatosensory cortices. The principal excitatory neurons in layers II/III have large axons that project to other telencephalic areas, such as the cortex and the striatum (Adesnik and Naka, 2018), while neurons in layer IV form loops within the layer and connect to layers II/III and VI (Scala et al., 2019). Layers VI and VII are thought to be the main outputs of the cerebral cortex, connecting to multiple subcortical areas and the thalamus, respectively (Harris and Shepherd, 2015). The higher C-FOS density in upper cortical layers indicates higher neuronal activity. This could reflect feedforward loops within neuronal populations and translaminar connectivity that could reinforce learning. However, further research is needed to determine the identity of these neurons.

Together, these findings give us a better understanding of the networks underlying eyeblink conditioning and provide candidate brain areas to be further researched in the context of associative learning.

## Materials and Methods

### Animals

All experiments were performed in accordance with the European Communities Council Directive. All animal protocols were approved by the Dutch National Experimental Animal Committee (DEC). C57BL/6J mice were ordered from Charles River (n = 16 males; n = 16 females), and B6CBAF1 mice from Janvier (n = 7 males; n = 9 females). Mice were group-housed and kept on a 12-hour light-dark cycle with ad libitum food and water. All procedures were performed in male and female mice approximately 8-12 weeks of age.

### Eyeblink pedestal placement surgery

Mice were anesthetized with isoflurane and oxygen (4% isoflurane for induction and 2-2.5% for maintenance). Body temperature was monitored during the procedure and maintained at 37°C. Animals were fixed in a stereotaxic device (Model 963, David Kopf Instruments, Tujunga CA, USA). The surgery followed previously described standard procedures for pedestal placement (Gao et al., 2016; ten Brinke et al., 2017). In short, the hair on top of the head was shaved, betadine and lidocaine were applied on the skin and an incision was done in the scalp to expose the skull. The tissue on top of the skull was removed and the skull was kept dry before applying Optibond™ prime adhesive (Kerr, Bioggio, Switzerland). A pedestal equipped with a magnet (weight ∼1g), was placed on top with Charisma^®^ (Heraeus Kulzer, Armonk NY, USA), which was hardened with UV light. Rymadil was injected subcutaneously (5mg per kg). Mice were left under a heating lamp for recovery for at least 3 hours. Mice were given 3-4 resting days before starting experiments.

### Eyeblink conditioning

Mice were habituated to the set-up (head fixed to a bar suspended over a cylindrical treadmill in a sound and light isolating chamber) for 5 days with increasing exposure (15, 15, 30, 45 and 60 min). Training started after two rest days. Twenty-eight 57BL/6 mice (n = 16 males; n = 16 females), and 16 B6CBAF1 mice (n = 7 males; n = 9 females) were trained using the standard eyeblink protocol (Brinke et al., 2015, Koekkoek et al., 2002). Ten CS-only trials of 30 ms with an inter-trial interval (ITI) of 10 ± 2 s were presented before the first training session to acquire a baseline measurement. Mice were next trained for 5 consecutive days. Each session consisted of 20 blocks of 12 trails each (1 US only, 11 paired and 1 CS only) with an ITI of 10 ± 2 s. The CS was a 270 ms blue LED light (∼ 450 nm) placed 7 cm in front of the mouse. The US was a 30 ms corneal air puff co-terminating with the CS. The puffer was controlled by a VHS P/P solenoid valve set at 30 psi (Lohm rate, 4750 Lohms; Internal volume, 30 μL, The Lee Company®, Westbrook, US) and delivered via a 27.5 mm gauge needle at 5 mm from the center of the left cornea. The inter-stimulus interval was 250 ms. Eyelid movements were recorded with a camera (Baseler aceA640) at 250 frames/s. 4 C57BL/6 mice (n = 2 males; n = 2 females), were trained using a pseudoconditioning protocol. Pseudoconditioning protocol consisted of 20 blocks of 12 trails each (1 puff only, 12 LED only) with an ITI of 10 ± 2 s. The puff and LED stimulus had the same characteristics as in the conditioning protocol. Data was analyzed with a custom written MATLAB code as previously described (Giovannucci et al., 2017; Badura et al., 2018). Traces were normalized within each session to the UR max amplitude. The CR detection window was set to 650-730 ms and CRs were only classified as such when the amplitude was equal or higher than 5% of the UR median. The CR percentage was calculated as the number of counted CRs (equal or higher than 5% of the UR median) divided by the total CS trials per session.

### Locomotion

An infrared camera (ELP 1080P) (sampling frequency 60 frames/s) was placed in each of the eyeblink boxes and connected to an external computer (independent from the eyeblink system). The cameras were positioned at the right back corner of the chamber on top of a magnet tripod attached to a custom-made metal block which allowed stable fixation. The recording angle was standardized by selecting the same reference in the field of view of each camera. Simultaneous video acquisition from the three cameras was performed in Ipi Recorder software (http://ipisoft.com/download/). Body movement recording was parallel to eyelid recording during the training sessions. The output videos (.avi format) from each mouse and session were approximately 35 min (corresponding to the length of an eyeblink session).

### Locomotion analysis

We used DeepLabCut (DLC) to track body parts from videos (Mathis et al., 2018) (**Fig. 2**). We extracted 40 frames of 4 different videos from two males and two females (total of 160 frames). Next, frames were manually labeled with 5 body parts (tail base, hip, knee, right back paw and nose). These frames were used for training the pre-trained deep neural network ResNet50 (He et al., 2016; Insafutdinov et al., 2016). Evaluation of the network was done to confirm a low error in pixels between labeled frames and predictions. Video analysis was done by using the trained network to get the locations of body parts from all mice and sessions (16 mice x 5 sessions = 80 videos). DLC output is a matrix with x and y positions in pixels and the likelihood of this position for each body part. We used this matrix to calculate distance covered and speed per body part with a custom written code (https://github.com/BaduraLab/DLC_analysis). After confirming normal distribution of the spatial coordinates per body part over training sessions, we performed the Grubbs’s test for outlier removal to discard possible tracking errors.

### Tissue processing

Mice were anesthetized with 0.2 ml pentobarbital (60 mg/ml) and perfused with 0.9% NaCl followed by 4% paraformaldehyde (PFA). Given the peak time expression of *C-fos* (Chung, 2015), animals were perfused 90 minutes after finishing the last training session. Brains were dissected from the skull and stored in 4% PFA at room temperature (rT) for 1.5 hours. They were next changed to a 10% sucrose solution and left overnight at 4°C. Brains were embedded in 12% gelatin and 10% sucrose and left in a solution with 30% sucrose and 4% PFA in PBS at rT for 1.5 hours. Next, they were transferred to a 30% sucrose solution in 0.1 M PB and kept at 4°C. Whole brains were sliced at 50 μm with a microtome and slices were kept in 0.1 M PB.

### Immunostaining and Imaging

Sections were incubated in blocking solution (10% NHS, 0.5% Triton in PBS) for an hour at rT. After rinsing, sections were incubated for 48 hours at 4°C on a shaker in primary antibody solution with 2% NHS (1:2000 Rabbit anti-C-FOS, ab208942, Abcam; 1:1000 Rat anti-Ctip2, ab18465, Abcam; 1:1000 Rabbit anti-GABAalpha6, G5555, Sigma-Aldrich; 1:1000 Rabbit anti-Cux1, (Ellis et al., 2001)). After rinsing, sections were incubated for 2 hours at rT on a shaker with secondary antibody (1:500 Donkey anti-rabbit A594, 711-585-152, Jackson; 1:500 Donkey anti-Rabbit A488, 711-545-152, Jackson; 1:500 Donkey anti-rabbit Cy5, 711-175-152, Jackson; Donkey anti-rat Cy3, 712-165-150, Jackson). Sections were counterstained with DAPI. Finally, sections were rinsed in 0.1 PB, placed with chroomulin on coverslips and mounted on slide glasses with Mowiol.

Sections were imaged with a Zeiss AxioImager 2 (Carl Zeiss, Jena, Germany) at 10x. A DsRed filter and an exposure time of 300 ms was used for the Alexa 595 channel (C-FOS). The DAPI channel was scanned at 20 ms or 30 ms exposure time. Tile scans were taken from whole brain slices. We processed half the sections obtained from slicing, hence, the distance between tile scan images was 100 μm. High resolution images were taken with a LSM 700 confocal microscope (Carl Zeiss, Jena, Germany).

### Image analysis

We developed an image analysis workflow for brain region identification and quantification of C-FOS positive neurons following eyeblink conditioning (**Supp. Fig. 1**) (https://github.com/BaduraLab/cell-counting). The workflow combines Fiji and a SHARP-Track, a software written in MATLAB initially developed to localize brain regions traversed by electrode tracks (Shamash et al., 2018) (https://github.com/cortex-lab/allenCCF/tree/master/SHARP-Track). Brain slices were preprocessed (rotating, cropping and scaling) with a custom written macro in Fiji (Schindelin et al., 2012). Next, slices were registered to the Allen Brain Atlas using the SHARP-Track user interface. Segmentation was performed on the registered slices in Fiji. Given the characteristic C-FOS staining pattern in the cerebellar granule layer (**Fig 3. A**), we used different thresholding algorithms for the cerebellum and for the rest of the brain. Following that, automated cell counting of C-FOS positive neurons was performed with a custom written macro in Fiji (cerebellum - circularity: 0.5-1, size: 0-20 pixels, rest of the brain - circularity: 0.7-1, size: 0-40 pixels) to get the X and Y coordinates of every detected cell. The output matrix of coordinates was used to create a ROI array per slice in SHARP-Track. This step allows one to one matching between the ROI array and the previously registered slice. Finally, the reference-space locations and brain regions of each neuron were obtained by overlapping the registration array with the ROI array. ROI counts were normalized by brain region surface following the hierarchical structure of the Allen Brain Atlas. The surface of each brain areas was calculated per slice and cell density was defined as ROI counts/surface.

### Statistics

Statistics were performed in MATLAB and GraphPad Prism 6. Data is reported as mean ± std or sem. Normality was tested and accepted for both eyeblink CR amplitudes and for speed of the right back paw. The corresponding statistical test for the *p* values reported are specified in Results. Time data (training sessions) was analyzed using two-way repeated measures ANOVA for sex and session. Sex effect is reported in Results, session effect is significant in all groups (indicating learning through time) and interaction is reported if significant. For Kendalls’s correlation on C-FOS data, we report Tau and *p* values.

## Acknowledgments

We thank Roxanne ter Haar, Elize D. Haasdijk and Stephanie Dijkhuizen for their help with experiments. This work was supported by Dutch Research Council (Vidi/ZonMw/917.18.380,2018).

