## Supplemental figures for "Eyeblink conditioning performance and brain-wide *C-fos* expression in male and female mice"

**Supplementary Information**

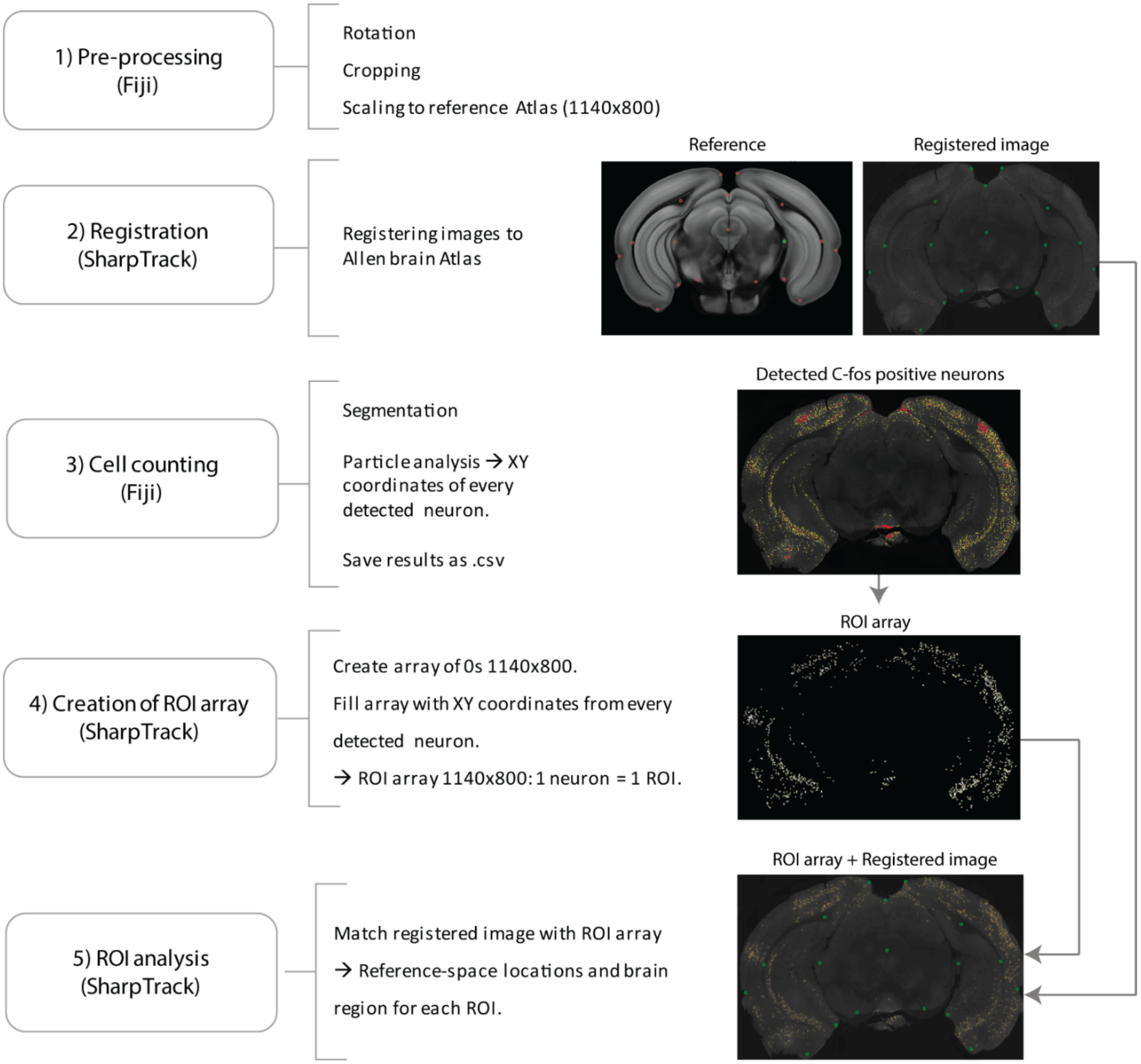


**Supplementary Figure 1:** Schematic analysis pipeline for C-FOS quantification.

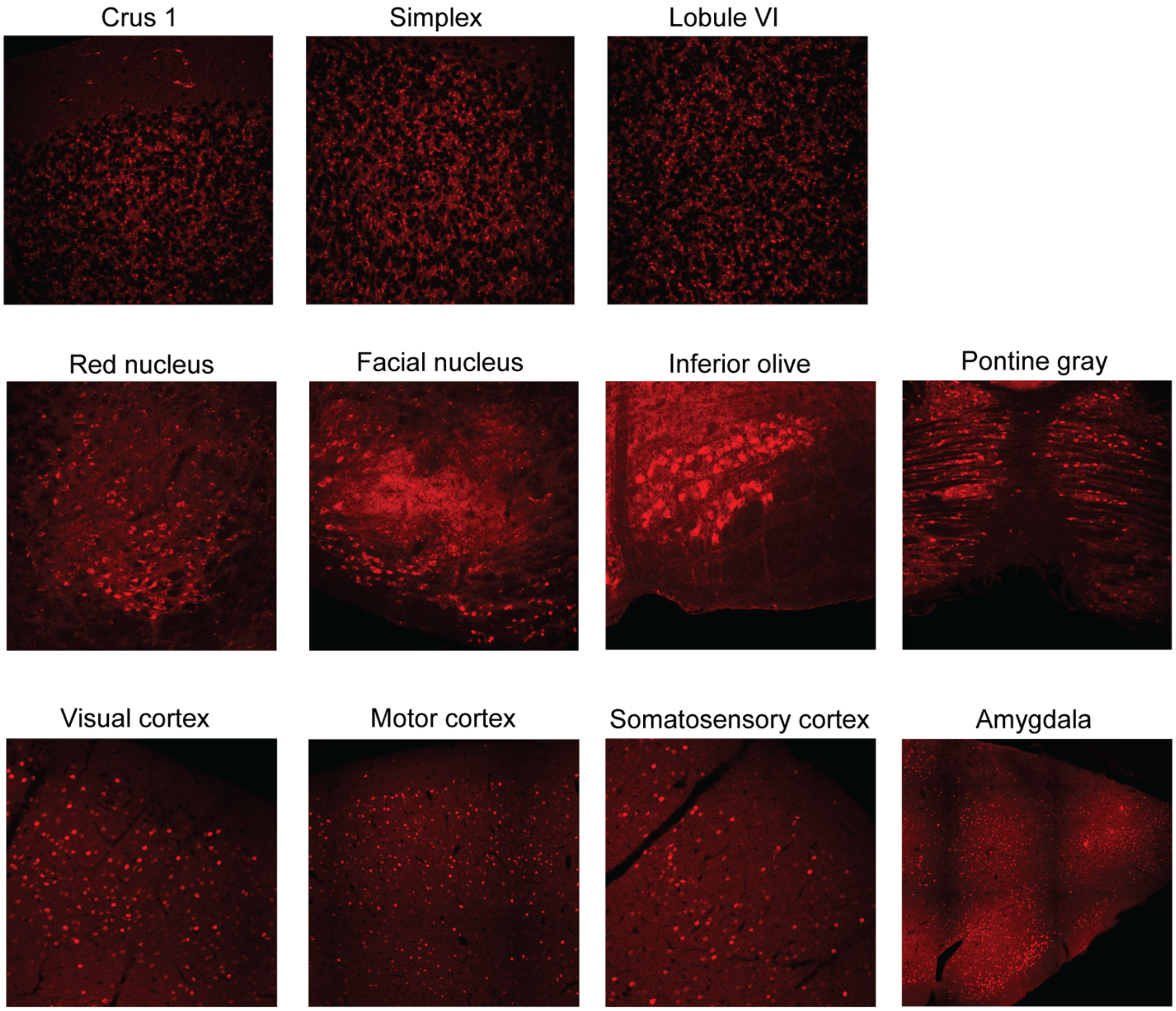


**Supplementary Figure 2:** Confocal images, brain areas that have a positive correlation between C-FOS cell density and CR amplitude in B6 mice. Cerebellum: 60X, others: 40X.

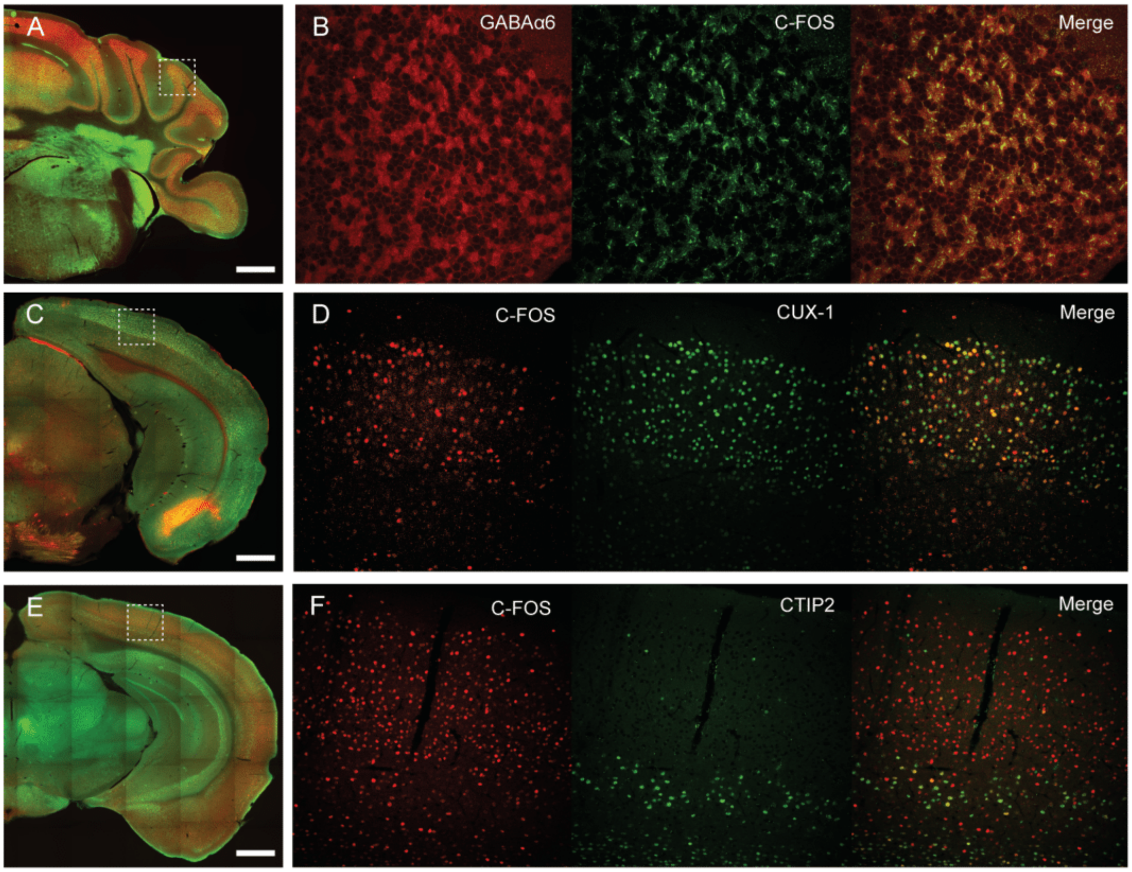


**Supplementary Figure 3:** Co-localization of C-FOS and other neuronal markers**. A**) crus 1, example image used for quantification of C-FOS positive cells. Red: GABAα6, Green: C-FOS. Scale: 500 µm. **B**) Confocal image of the zoomed in area in A, 60x. **C**) visual cortex, example image used for quantification of C-FOS positive cells. Red: C-FOS, green: CUX-1. Scale: 500 µm. **D**) Confocal image of the zoomed in area in C, 40X. **E**) visual cortex, example image used for quantification of C-FOS positive cells. Red: C-FOS, green: CTIP2. Scale: 500 µm. **F**) Confocal image of the zoomed in areas in E, 40X.

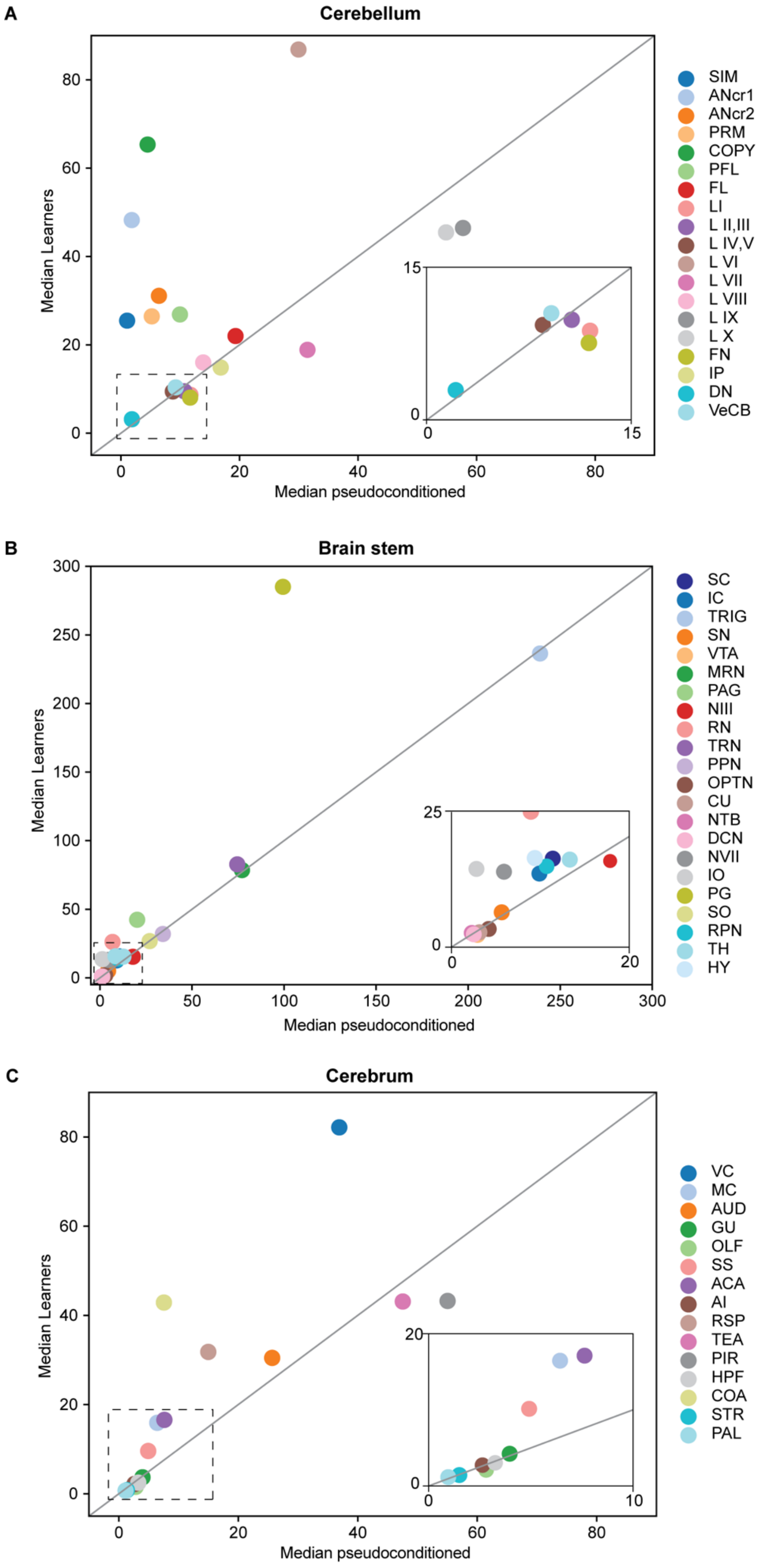
A
Cerebellum
Median Learners
Median pseudoconditioned
SIM
ANcr1
ANcr2
PRM
COPY
PFL
FL
LI
L II,III
L IV,V
L VI
L VII
L VIII
L IX
L X
FN
IP
DN
VeCB
B
Brain stem
SC
IC
TRIG
SN
VTA
MRN
PAG
NIII
RN
TRN
PPN
OPTN
CU
NTB
DCN
NVII
IO
PG
SO
RPN
TH
HY
C
Cerebrum
VC
MC
AUD
GU
OLF
SS
ACA
AI
RSP
TEA
PIR
HPF
COA
STR
PAL

**Supplementary Figure 4**: Median pseudoconditioned mice against median learners (CR amplitude session 5 > 0.4. Units in both axis are C-FOS positive cells/mm2. Pseudoconditioned mice: n = 2 B6 males, 2 B6 females. Learners n = 5 B6 males, 9 B6 females. The abreviations shown in the graph correspond to the Allen Brain Atlas abreviations for brain areas. Cerebellum: Hemispheric regions: SIM: simplex, ANcr1: crus 1, ANcr2: crus 2, PRM: paramedian, COPY: copula pyramidis, PFL: paraflocculus, FL: flocculus, Vermal Regions: Lobules I to X, Cerebellar nuclei: FN: fastigial, IP: interposed, DN: dentate, VeCB: vestibulocerebellar. Brain stem: SC: superior colliculus, IC: inferior colliculus, TRIG: trigeminal nucleus, SN: substantia nigra, VTA: ventral tegmental area, MRN: midbrain reticular nucleus, PAG: periaqueductal grey, NIII: oculomotor nucleus, RN: red nucleus, TRN: tegmental nucleus, PPN: pedunculopontine nucleus, OPTN: optic tract nucleus, CU: cuneate nucleus, NTB: nucleus of the trapezoid body, DCN: dorsal column nucleus, NVII: facial nucleus, IO: inferior olive, PG: pontine nucleus, SO: superior olive, RPN: raphe nucleus, TH: thalamus, HY: hypothalamus. Cerebrum: VC: visual cortex, MC: motor cortex, AUD: auditory cortex, GU: gustatory cortex, OLF: olfactory cortex, SS: somatosensory cortex, ACA: anterior cingulate cortex, AI: insula, RSP: retrosplenial area, TEA: temporal association area, PIR: piriform area, HPF: hippocampus, COA: amygdala, STR: striatum, PAL: pallidum.

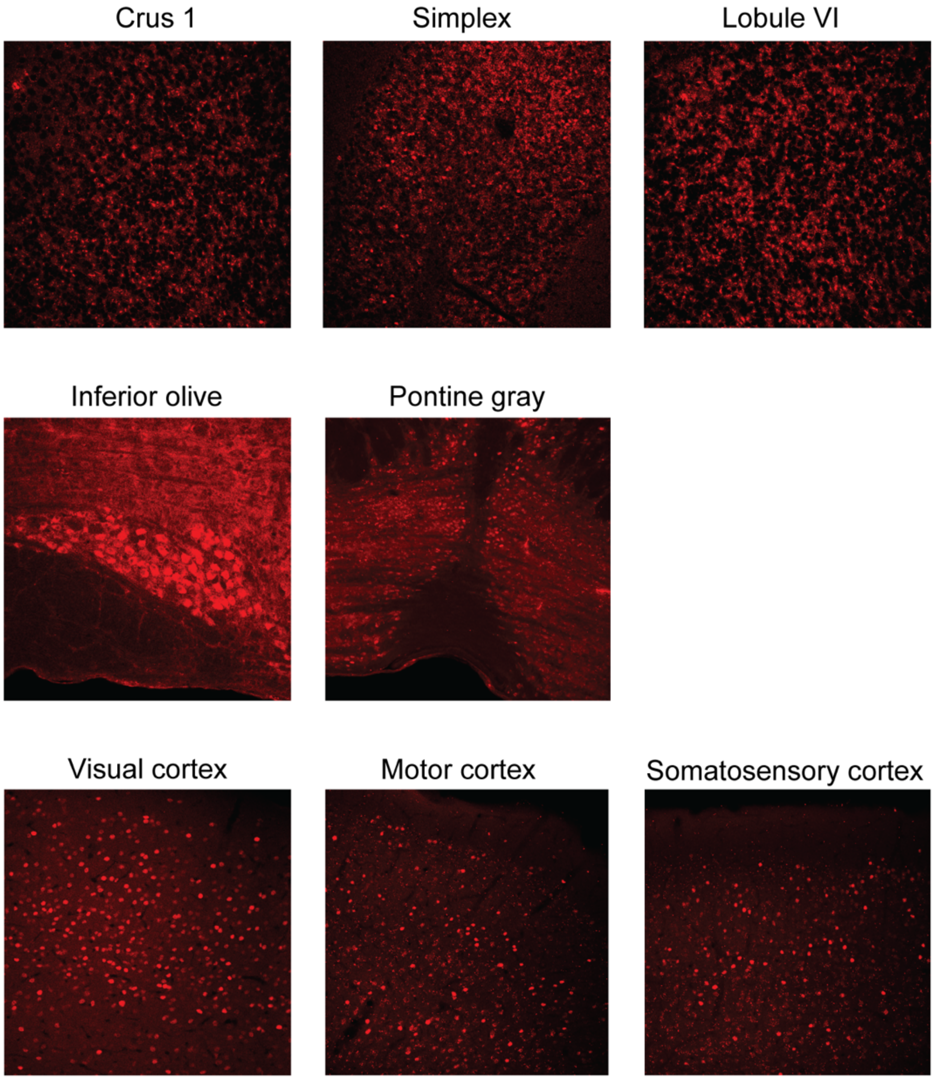


**Supplementary Figure 5**: Confocal images, brain areas that have a positive correlation between C-FOS cell density and CR amplitude in B6CBAF1 mice. Cerebellum: 60X, others: 40X.

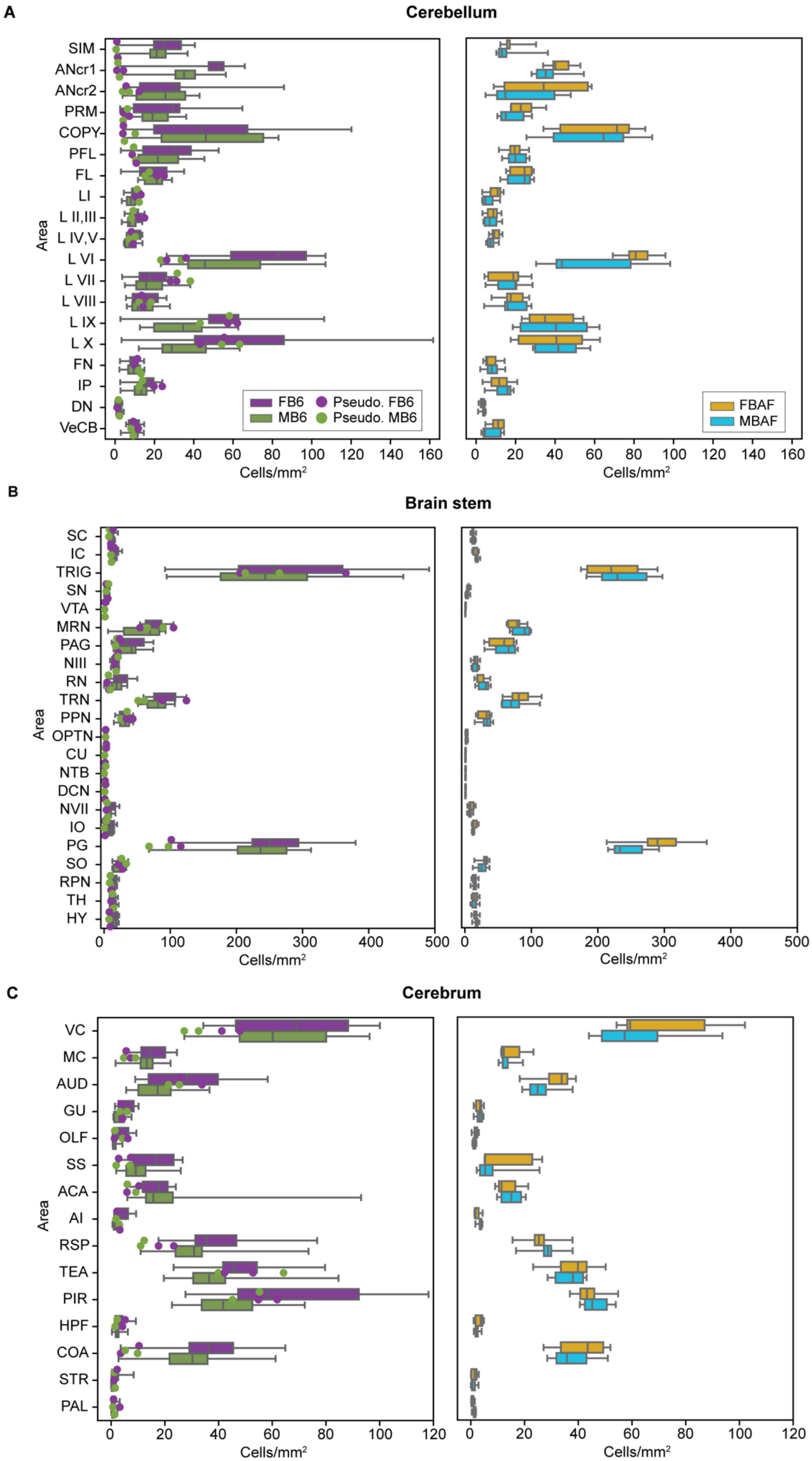
A
Cerebellum
SIM
ANcr1
ANcr2
PRM
COPY
PFL
FL
LI
L II,III
L IV,V
L VI
L VII
L VIII
L IX
L X
FN
IP
DN
VeCB
Area
FB6
MB6
Pseudo. FB6
Pseudo. MB6
FBAF
MBAF
Cells/mm²
B
Brain stem
SC
IC
TRIG
SN
VTA
MRN
PAG
NIII
RN
TRN
PPN
OPTN
CU
NTB
DCN
NVII
IO
PG
SO
RPN
TH
HY
C
Cerebrum
VC
MC
AUD
GU
OLF
SS
ACA
AI
RSP
TEA
PIR
HPF
COA
STR
PAL

**Supplementary Figure 6:** Boxplots showing the distribution of C-FOS positive cell density throughout the brain in all mice. B6 mice: conditioned animals: 14 males and 14 females, pseudoconditioned animals (dots): 2 males and 2 females. B6CBAF1 mice: 7 males and 9 females. The abreviations shown in the graph correspond to the Allen Brain Atlas abreviations for brain areas. Cerebellum: Hemispheric regions: SIM: simplex, ANcr1: crus 1, ANcr2: crus 2, PRM: paramedian, COPY: copula pyramidis, PFL: paraflocculus, FL: flocculus, Vermal Regions: Lobules I to X, Cerebellar nuclei: FN: fastigial, IP: interposed, DN: dentate, VeCB: vestibulocerebellar. Brain stem: SC: superior colliculus, IC: inferior colliculus, TRIG: trigeminal nucleus, SN: substantia nigra, VTA: ventral tegmental area, MRN: midbrain reticular nucleus, PAG: periaqueductal grey, NIII: oculomotor nucleus, RN: red nucleus, TRN: tegmental nucleus, PPN: pedunculopontine nucleus, OPTN: optic tract nucleus, CU: cuneate nucleus, NTB: nucleus of the trapezoid body, DCN: dorsal column nucleus, NVII: facial nucleus, IO: inferior olive, PG: pontine nucleus, SO: superior olive, RPN: raphe nucleus, TH: thalamus, HY: hypothalamus. Cerebrum: VC: visual cortex, MC: motor cortex, AUD: auditory cortex, GU: gustatory cortex, OLF: olfactory cortex, SS: somatosensory cortex, ACA: anterior cingulate cortex, AI: insula, RSP: retrosplenial area, TEA: temporal association area, PIR: piriform area, HPF: hippocampus, COA: amygdala, STR: striatum, PAL: pallidum.
